# The Structural History of Eukarya

**DOI:** 10.64898/2026.02.04.703859

**Authors:** Qiuzhen Li, Diandra Daumiller, Patrick Bryant

**Affiliations:** Science for Life Laboratory, The Department of Molecular Biosciences, The Wenner-Gren Institute, Stockholm University, Solna 171 65, Sweden

**Author notes:** Equal contribution (Qiuzhen Li, Diandra Daumiller).

## Abstract

Comparative genomics has traditionally relied on sequence-based markers, yet the global evolutionary landscape of protein architecture remains largely unexplored. We present the Structural History of Eukarya (SHE), a phylogeny derived from all-vs-all structural comparisons of 1,542 eukaryotic proteomes, encompassing nearly 300 trillion protein-protein alignments. This proteome-scale analysis reveals a bipartite model of eukaryotic evolution: a rigid Strict Core dominated by cytoskeletal architecture, supporting a more plastic Operational Engine centred on translational machinery. We identify lineage-specific accelerations of structural evolution in species such as birds and ants that are decoupled from proteome expansion, and show that aggregate structural topology provides a quantitative diagnostic of reference proteome quality. Finally, we demonstrate that proteome-wide structural fidelity enables a data-driven framework for model organism selection, replacing heuristic choices with quantitative matching to human biological processes. SHE is freely accessible as an interactive portal: https://she-app.serve.scilifelab.se/

## Introduction

Historically, reconstructing the Tree of Life has relied on comparing a select group of highly conserved molecular markers, such as ribosomal RNA (rRNA) [1]. While this approach yields high-confidence phylogenetic trees for closely related organisms, its utility is constrained by the omission of vast evolutionary data contained within the broader genome. This reliance is further complicated by heterotachy, varying evolutionary rates across different genes and lineages, which introduces significant uncertainty as the divergence between organisms increases. Such limitations in sequence-based phylogeny do more than obscure ancestral relationships; they compromise the reliability of critical applications, including phylogenomic-based gene function annotations [2].

To overcome these hurdles, the three-dimensional (3D) structure of proteins has emerged as a more robust indicator of deep evolutionary history. Protein architecture is remarkably stable; because the biophysical requirements for folding are so stringent, structural cores evolve 3 to 10 times more slowly than their primary amino acid sequences [3]. This stability makes structural phylogenetics a powerful tool for resolving ancient evolutionary splits that sequence data can no longer accurately track [4]. Recent research demonstrates that structure-based models consistently outperform sequence-only methods across all levels of divergence, from distantly related proteins to those more closely aligned [5]. The versatility of this approach has already been validated through the successful reclassification of rapidly evolving viruses, highlighting its potential to redefine our understanding of the eukaryotic landscape [6].

Despite its theoretical advantages, large-scale structural phylogenetics has historically been bottlenecked by two formidable challenges: a scarcity of experimentally determined structures and the prohibitive computational cost of structural alignment [7][8]. For decades, the structural universe remained largely inaccessible, as the vast majority of proteins lacked high-resolution models, and the resources required to compare those that did were immense. However, a recent technological shift has effectively dismantled these barriers.

The emergence of high-accuracy AI models, most notably AlphaFold2 [9], has democratised structural biology by generating hundreds of millions of high-quality protein architectures, thereby populating the AlphaFold Protein Structure Database (AFDB) and resolving the long-standing data deficit [10].

Complementing this influx of data, FoldSeek has circumvented the computational burden of alignment by translating complex 3D geometries into a simplified "3Di" sequence alphabet [11]. This innovation transforms resource-intensive structural comparisons into rapid, sequence-like alignments without sacrificing sensitivity. Finally, the logistical challenge of managing these massive datasets has been addressed by Foldcomp, a high-performance compression format that allows for the efficient storage and rapid retrieval of millions of structures [12]. Together, these advancements have transitioned structural phylogenetics from a niche speciality into a scalable, proteome-wide methodology.

Harnessing these recent breakthroughs in structure prediction and high-throughput alignment, we present an all-against-all structural analysis of complete eukaryotic proteomes. Our study establishes a robust metric of proteome similarity that accounts for both the structural conservation of individual protein folds and variations in overall proteome size. This metric allows us to construct the Structural History of Eukarya (SHE), the most comprehensive and high-resolution eukaryotic phylogeny derived from structural data to date. Beyond reconstructing ancestral lineages, SHE introduces a novel framework for quantifying the rate of structural proteome evolution across the eukaryotic tree. By providing the complete dataset as an interactive, open-access resource, we offer an unprecedented lens through which to examine the evolutionary forces that have sculpted the diversity of eukaryotic life.

## Results

### The structural similarity of eukaryotic proteomes

Protein structures are generally more conserved than sequences since maintaining the correct 3D fold is crucial for function, while different amino acid sequences can achieve the same fold through compensatory mutations [3,13]. In addition, comparing organisms on the level of a single gene or protein results in the exclusion of all other proteome information. To obtain a comparison between organisms on the structural proteome level, we aligned all structures from the AlphaFold database [10] in all complete eukaryotic proteomes from UniProt (n=1542) [14] against each other using FoldSeek (1.2 million proteome alignments, ∼300 trillion protein-protein alignments between 24.3 million unique proteins). This constitutes the largest protein structural comparison to date (to our knowledge), and required vast computational resources and novel, highly efficient pipelines combining the FoldComp-MMseqs-FoldSeek ecosystem [11,12,15,16]. From the structural hits, we define the proteome similarity as

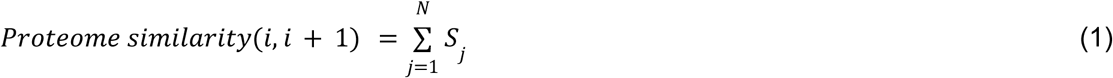

S_j_ is the highest FoldSeek structural bit score [11] (the top hit) obtained from searching protein j from proteome(i), consisting of proteins j..N, against proteome(i+1). We use each hit protein only once. The result is that proteomes of different sizes are penalised, and large proteomes with high similarities are favoured, as these will have more support. We define the proteome distance as the inverse of the similarity:

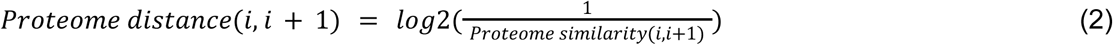

The resulting distances were used to construct the evolutionary history of protein structures across eukaryotic proteomes (**Figure 1, Methods**). The tree is the most complete and sensitive reconstruction to date of the protein structural history of Eukarya, and enables studies of evolutionary rates across proteomes, proteome-wide structural comparisons and the possibility to investigate the evolution of key cellular functions in a more complete context, all of which we present below. Importantly, all alignment scores are precomputed and stored, providing a valuable resource for efficient structural comparison across proteomes without expensive re-computation of pairwise relationships on any level.

**Figure 1.**
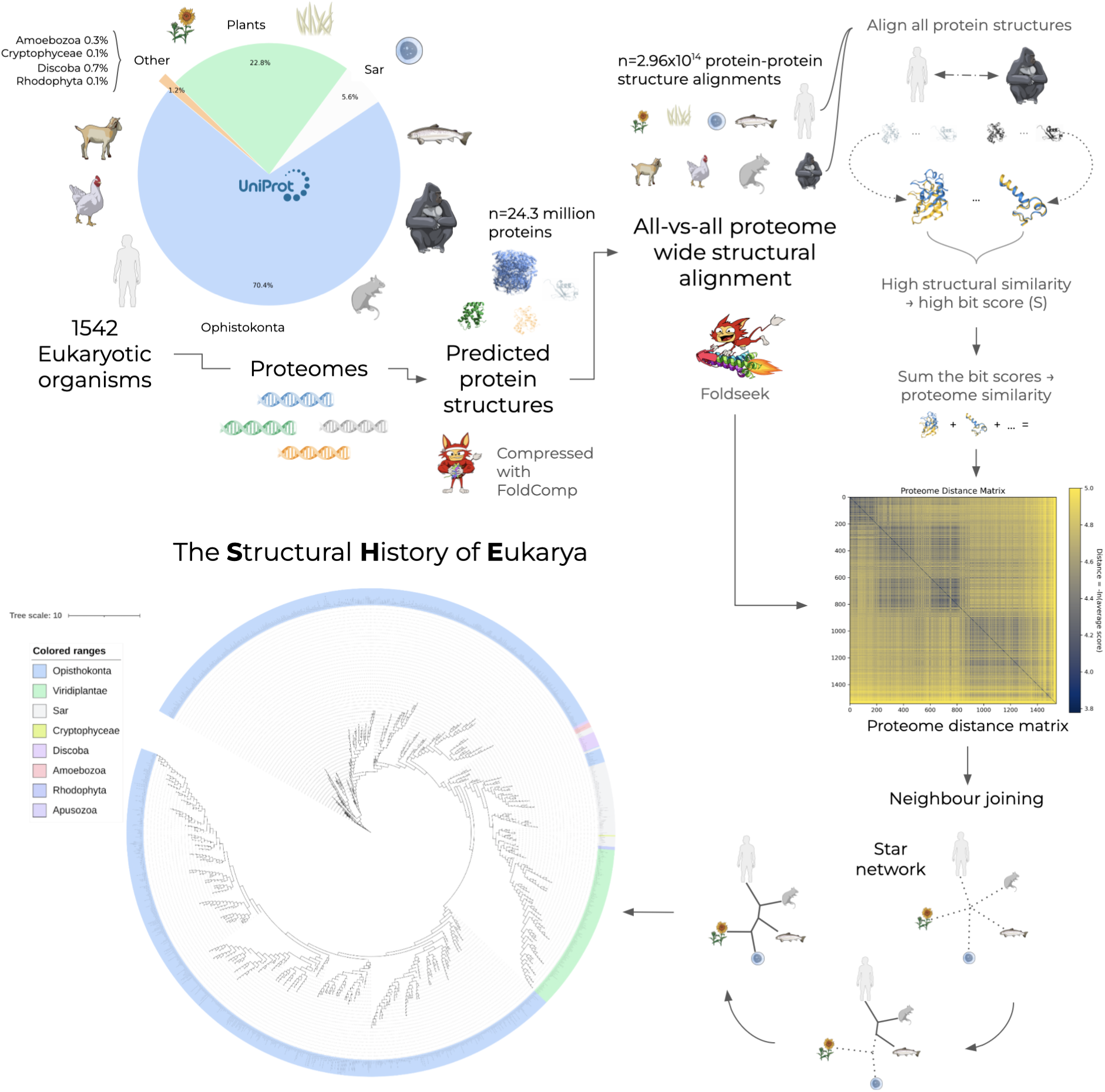
The Structural History of Eukaryotic Proteomes: a proteome-wide structural analysis workflow. To reconstruct evolutionary relationships at the structural level, we developed a high-throughput pipeline starting with the extraction of all eukaryotic proteomes from UniProt annotated as highly complete (median gene retention 96%). Predicted structures for these proteomes were retrieved from the AlphaFold Database (AFDB) and compressed using FoldCompto build proteome-specific structural databases. We then performed an all-against-all structural alignment using Foldseek, comparing every protein in each proteome against every protein in all others (exemplified by the human vs. gorilla comparison). This resulted in 1,188,111 proteome-level alignments comprising 2.96 × 10^14^ protein-protein alignments across 24,333,776 unique proteins. A Proteome Distance Matrix was generated by aggregating the highest-scoring bit scores for each protein pair (using the inverse bit score sum, Equation 2). Finally, a eukaryotic phylogenetic tree was constructed from this matrix using neighbor-joining with bootstrapping. The circular dendrogram visualizes these relationships, with major taxonomic groups highlighted: Opisthokonta (blue), Viridiplantae (green), SAR (Stramenopiles, Alveolates, Rhizaria; pink), Discoba (rose), Amoebozoa (purple), Rhodophyta (indigo), Cryptophyceae (lime green), and Apusozoa (violet). A scale bar indicating evolutionary distance (derived from Equation 2) is provided for reference.

### The Structural Evolution of Protein Sets

#### Data-Driven Model Organism Selection

SHE contains nearly 300 trillion precomputed protein-protein alignments between 24.3 million unique proteins. These are “ready-made”, which enables rapid comparison of sets of proteins for e.g. pathway comparison across Eukarya. To showcase the utility of such comparisons, take the example of finding a suitable model organism for studying a specific human signalling pathway or biochemical process. First, all human proteins have to be mapped to all possible proteins in all possible model organisms considered. Then, the results have to be compared across each pathway and potential species hit to collect support for the best match. With SHE, this is already computed. One can simply select a set of proteins in an organism among the 1542 studied, and either list a set of potential model organisms or search all. Then, the alignment scores can be extracted from the precomputed results and used to calculate a matching score across the protein set.

#### Case Study: The Fanconi Anaemia Pathway

As an example, we provide a quantitative ranking of model organisms for the study of the Fanconi Anaemia (FA) pathway (**Figure 2**). The query included key players in the pathway, FANCD2 and FANCI, which form a central hub to recruit downstream factors to the DNA interstrand crosslink (ICL) site [17], as well as the regulatory kinase CHK1 and the E3 ligase scaffold CUL1, which work with FBXL12 to mediate the degradation of FANCD2 during oncogene-induced replication stress [18]. As expected, the pathway architecture is exceptionally well-conserved in Mus musculus (bit score = 25,999), reinforcing its status as the primary mammalian model for FA research. In contrast, pathway scores decline sharply in non-vertebrate models; Drosophila melanogaster and Saccharomyces cerevisiae yielded significantly lower scores (2790 and 341, respectively). These computational results align closely with established biological literature, as the canonical FA pathway is known to be absent or highly divergent in flies and yeast, which utilise alternative ICL repair mechanisms [19]. Interestingly, while the vertebrate Danio rerio (bit score = 7063) retains a functional version of the pathway, Caenorhabditis elegans (bit score = 7094) exhibits a comparable score, suggesting its potential as a streamlined model for investigating specific structural components of the FA machinery.

**Figure 2:**
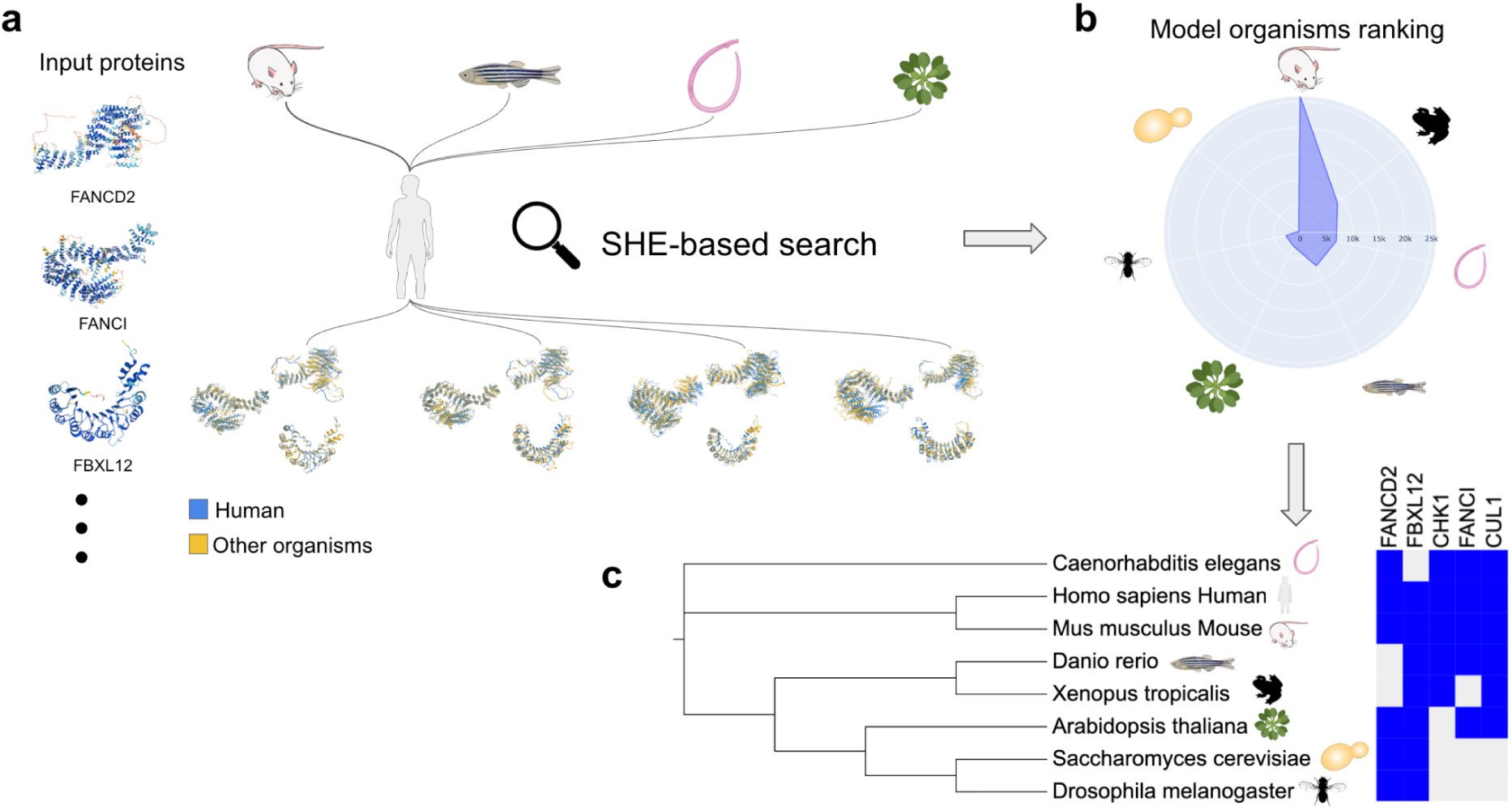
Quantitative ranking of model organisms for the Fanconi Anaemia pathway. **a)** Schematic workflow of the SHE-based (Sequence Homology and Evolutionary) search strategy. Key human FA pathway proteins (FANCD2, FANCI, FBXL12, CHK1 and CUL1) were used as query inputs to identify orthologs across diverse species. **b)** Model organism ranking based on cumulative bit scores. *Mus musculus* ranks highest, indicating strong pathway conservation, whereas non-vertebrates show lower scores. **c)** Phylogenetic clustering and protein conservation heatmap. The heatmap indicates the presence (blue) or absence (grey) of specific FA components.

#### Family-specific Phylogeny

Although SHE is designed as a proteome-level resource, its architecture allows for the instantaneous extraction of family-specific phylogenies without the need for *de novo* sequence alignment. By "pruning" the global species tree to retain only those lineages possessing a specific structural homolog, the platform allows users to visualise the evolutionary trajectory of distinct protein families in real time.

To demonstrate this capability, we contrasted the phylogenetic footprints of two functionally distinct families (**Figure 3**). The Sodium/potassium-transporting ATPase subunit alpha (ATP1A) family exhibits a ubiquitous distribution across the analysed lineages, yielding a dense, broadly branched sub-tree consistent with its essential "housekeeping" role in animal physiology [20]. In contrast, the FXNA-like protease family reveals a highly restricted distribution, found exclusively within the *Drosophila* clade. Because SHE relies on pre-computed structural data, these pruned topologies are generated instantly, enabling researchers to efficiently distinguish between ancient, universally conserved machineries and those that have evolved for specialised, taxon-specific roles.

**Figure 3.**
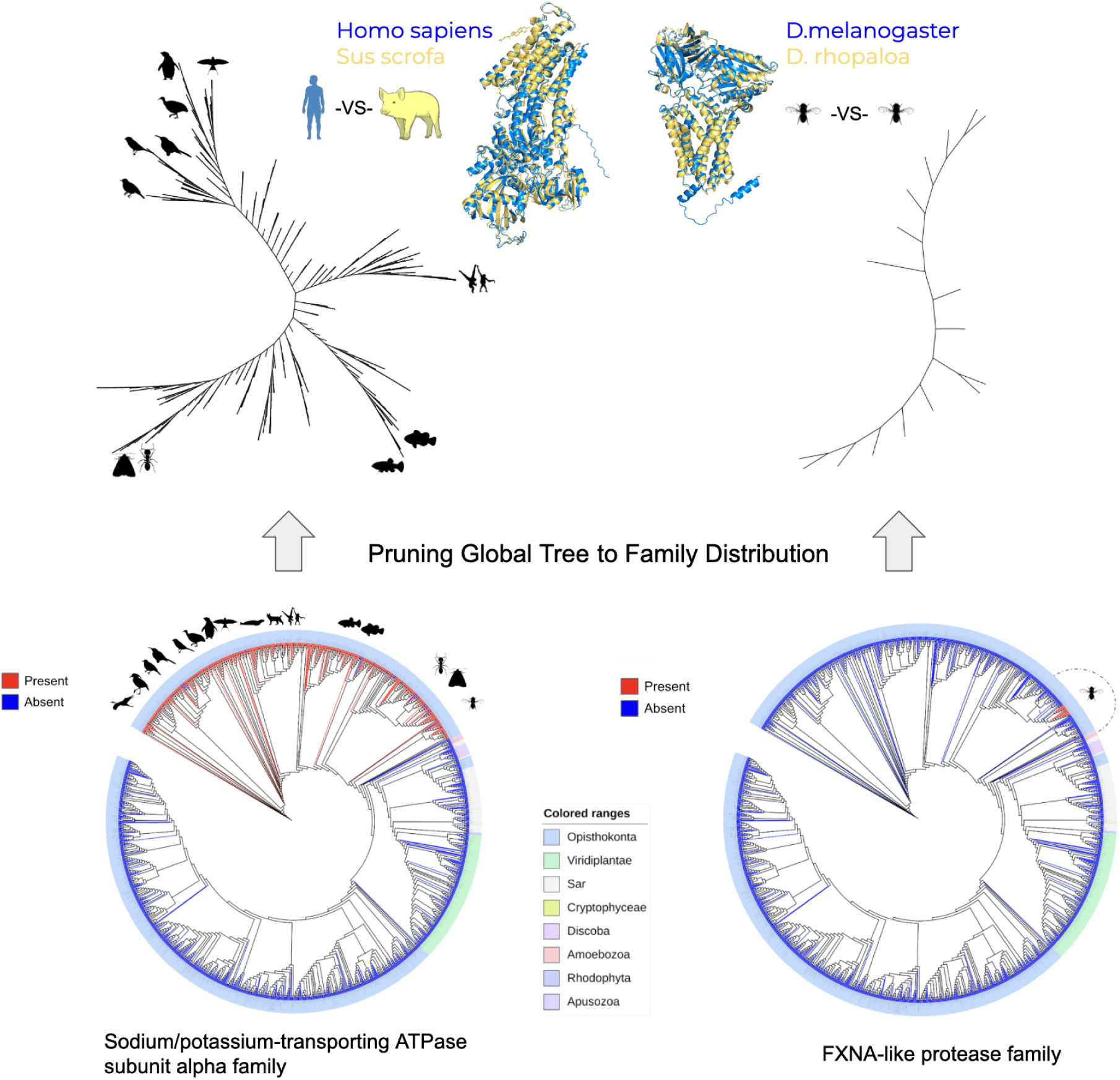
SHE: A solution for the rapid analysis of homologous protein families. The platform enables the seamless transition from global phylogenetic mapping to high-resolution clade analysis. **Bottom panels:** The global phylogenetic distribution of two distinct protein families is mapped across the eukaryotic tree, where red branches indicate the presence of a structural homolog and blue branches indicate absence. The outer ring provides a taxonomic guide to eukaryotic kingdoms according to the colored legend. **Top panels:** Extracted sub-trees illustrate the corresponding evolutionary relationships within the retained species. **(Left)** The ATP1A family (Sodium/potassium-transporting ATPase subunit alpha) exhibits a broad, conserved distribution across *Opisthokonta*, yielding a dense extracted phylogeny characteristic of an essential "housekeeping" protein. The accompanying structure shows a superposition of homologs, highlighting the conservation between homo sapiens and Sus scrofa. **(Right)** In contrast, the FXNA-like protease family reveals a highly restricted, lineage-specific distribution found exclusively within the *Drosophila* clade. The extracted sub-tree and structural alignment focus on specific relationships, shown here as the comparison between D. melanogaster and D. *rhopaloa*.

### Evolutionary Rates Across Proteomes

SHE enables the precise analysis of evolutionary rates of structural change across eukaryotic lineages. We find that the tempo of structural proteome evolution is highly variable across kingdoms and branching points. To quantify these dynamics, we defined the rate of proteome change as:

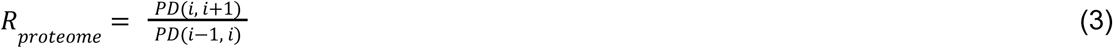

Where PD(i, i+1) is the proteome distance of proteomes i and i+1 and PD(i-1, i) that of proteomes i-1 and i in the order of the obtained phylogeny directed towards the edge nodes. An R_proteome_ > 1 indicates an acceleration in structural divergence, while R_proteome_ < 1 indicates a deceleration.

When examined at the kingdom level, the distributions of these rates show significant variation (**Figures 4b&c**). While baseline rates generally remain low across most lineages, Opisthokonta, SAR (Stramenopiles, Alveolates, Rhizaria), and Viridiplantae all exhibit extreme outliers with change rates reaching the thousands, suggesting periods of rapid evolutionary restructuring within relatively short time frames. (Note: The Apusozoa and Cryptophyceae lineages were represented by a single species, precluding distribution analysis). Such variation is a well-known phenomenon in molecular evolution, often linked to biological factors like generation time or metabolic rate [21], but has not previously been characterised at the global proteome level.

**Figure 4.**
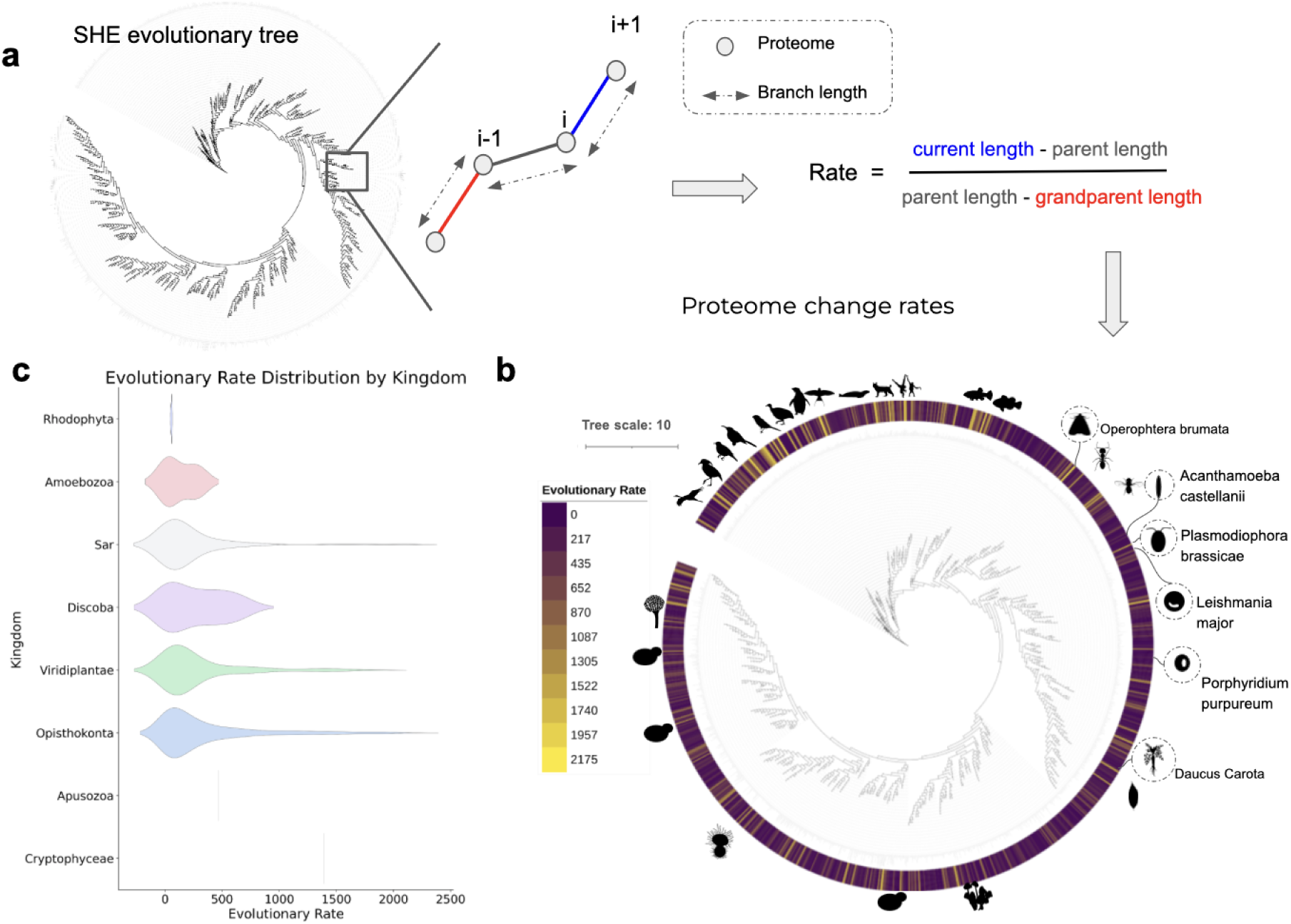
Evolutionary rate measured as the rate of proteome structural change. **a)** Schematic of the evolutionary rate calculation. The rate of evolution for a given branch is quantified as the ratio of branch lengths between successive nodes (i-1, i, i+1). As defined in Equation 3, this metric compares the magnitude of the most recent structural divergence (blue segment) against the preceding interval (red segment), providing a measure of evolutionary acceleration (R > 1) or deceleration (R < 1). **b)** Global phylogenetic tree of evolutionary rates. The eukaryotic tree is colored according to the calculated rate of structural change, ranging from slow (dark magenta) to fast (yellow). Silhouettes highlight species with exceptionally high rates (>1,500), with the top-ranking outlier labelled for each major clade (e.g., *Operophtera brumata* for Opisthokonta, *Daucus carota* for Viridiplantae). Note the distinct cluster of accelerated evolution observed within the Class Aves (birds). **c)** Distribution of evolutionary rates by kingdom. Violin plots visualise the density of evolutionary rates across the major eukaryotic supergroups: Opisthokonta (blue), Viridiplantae (green), SAR (pink), Discoba (rose), Amoebozoa(purple), Rhodophyta (indigo), Cryptophyceae (lime green), and Apusozoa (violet). Wider sections of the violin represent a higher frequency of species at that specific rate.

The highest rates of structural evolution identified within each major clade are summarised in **Table 1**. Within Opisthokonta, we observed notably accelerated evolution in Aves (birds), Primates, Teleostei (bony fishes), and Formicidae (ants) (**Figure 4b**). This finding, derived entirely from our structural phylogenetic approach, corroborates the rapid evolutionary rates previously reported for groups like *Neoaves* using genomic methods [15]. Crucially, this variation in evolutionary speed appears to be intrinsic to the lineage rather than a function of data magnitude. We observed no significant correlation between structural conservation and proteome size (Spearman’s ρ = 0.04) (**Figure 5a**). This result challenges simplistic models that link proteome complexity to evolutionary rate [22], suggesting that structural innovation is driven by distinct evolutionary pressures.

**Figure 5.**
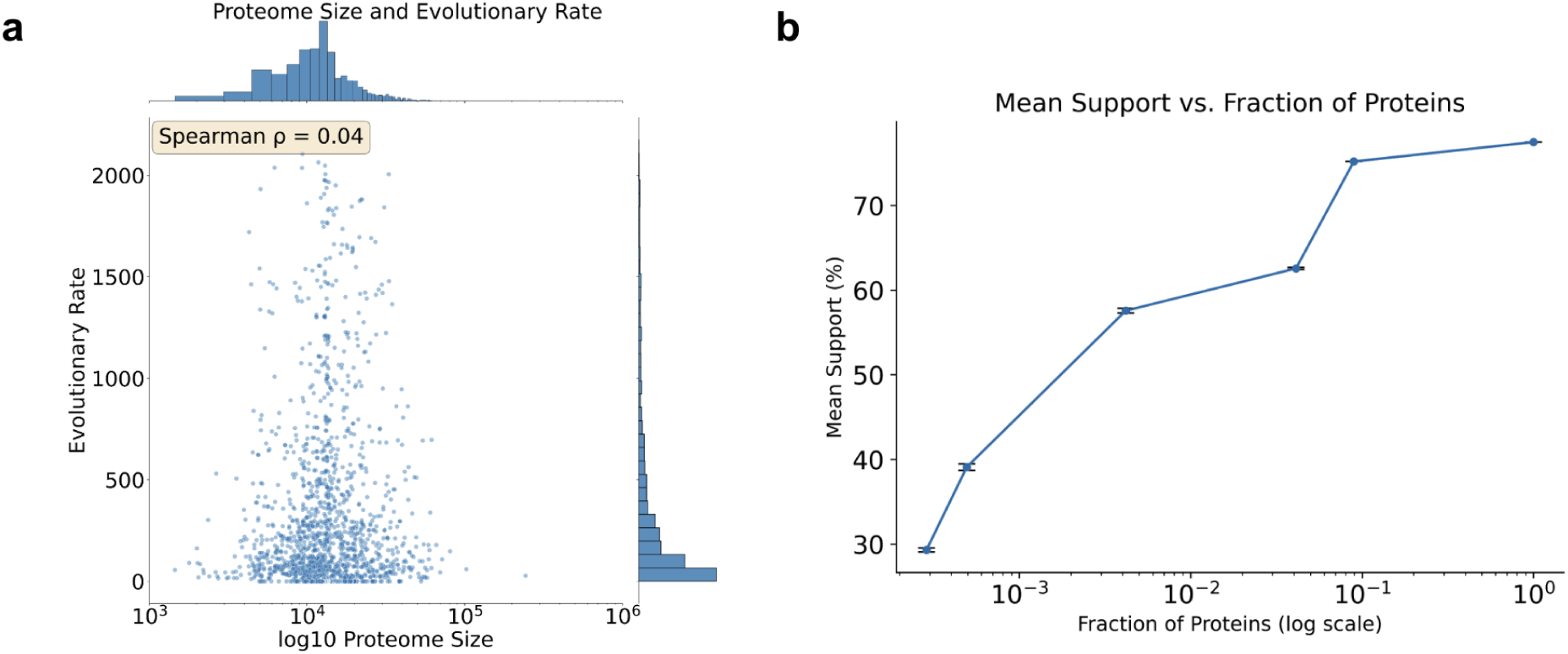
The effect of proteome size and sampling depth on evolutionary rates and phylogeny. **a)** Correlation between proteome size and evolutionary rate. A scatter plot comparing total proteome size (log scale) against the calculated evolutionary rate (R_proteome_) for each species. Marginal histograms display the distribution of each variable. The Spearman rank correlation coefficient (ρ = 0.04) indicates no significant relationship, suggesting that structural evolutionary rates are independent of proteome size. **b)** Impact of protein subset size on phylogenetic support. The plot illustrates the relationship between the fraction of the total proteome included in the analysis (x-axis, log scale) and the mean bootstrap support for the resulting phylogenetic tree (y-axis). Data points represent the average of five independent runs with standard deviation bars. The trend confirms that increasing the volume of structural data significantly enhances the statistical robustness of the inferred phylogeny.

**Table 1.** Maximum rates of structural proteome evolution across major eukaryotic clades. The table lists the species exhibiting the highest calculated rate of structural change (R_proteome_) within their respective supergroups. The rate is derived from Equation 3 and represents the ratio of structural divergence between successive nodes in the phylogenetic tree, where higher values indicate accelerated evolutionary shifts. Common names are provided where applicable.

| Clade | Species | Common Name | $R_{proteome}$ |
| --- | --- | --- | --- |
| Opisthokonta | *Operophtera brumata* | Winter Moth | 2,175.3 |
| SAR | *Plasmodiophora brassicae* | Clubroot | 2,105.3 |
| Viridiplantae | Daucus carota | Carrot | 1,842.4 |
| Discoba | Leishmania major | - | 669.1 |
| Amoebozoa | Acanthamoeba castellanii | - | 277.0 |
| Rhodophyta | Porphyridium purpureum | - | 57.97 |

### Proteome-Scale Data Enhances Phylogenetic Resolution

A central premise of the SHE framework is that integrating structural information from complete proteomes offers superior phylogenetic resolution compared to single-gene markers. To validate this, we systematically measured how phylogenetic resolution scales with the volume of structural data included. We generated five distinct distance matrices based on increasingly larger subsets of the most common protein families, ranging from 7,000 to 2.1 million proteins. These were compared against the master phylogeny derived from the full dataset of 24.3 million proteins. For each subset, we constructed a neighbour-joining (NJ) tree and evaluated statistical robustness using 10,000 bootstrap replicates (see Methods).

Our analysis reveals a strong positive correlation between data volume and the statistical support for the tree’s topology (**Figure 5b**). A phylogeny based on a single protein family (∼7,000 proteins) yielded a mean bootstrap support of only 29.6%. In this limited tree, only 55 of 317 internal nodes (17%) achieved strong branching support (≥ 70 %). As expected, increasing the structural coverage dramatically improved resolution. Mean bootstrap support rose to 39.5% with ∼12,000 proteins and reached 57.2% with a subset of 100,000. This trend continued upward, culminating in a mean bootstrap support of 77.4% for the tree constructed from the full 24.3 million protein dataset. These results demonstrate that the aggregation of proteome-wide structural signals reliably and significantly strengthens inferred evolutionary relationships.

### The Core of Eukaryotic Proteomes

Identifying the proteins that are consistent across all eukaryotes is fundamental to reconstructing the biology of the Last Eukaryotic Common Ancestor (LECA) [23]. Traditionally, studies aiming to define the LECA proteome have relied on sequence-based methods to identify Eukaryotic Orthologous Groups. These analyses established that the ancestor was a complex cell, possessing a core proteome of several hundred to a few thousand proteins dedicated to information processing, membrane trafficking, and metabolism [24].

With SHE, we can refine this view by identifying the "structural core"—a subset of proteins that maintain their 3D fold across the entire eukaryotic tree, independent of sequence divergence. We established orthologous families using MCL clustering (see Methods) and filtered for those present across the vast majority of lineages. We specifically examined the ’Strict Core’ proteome, defined as protein families conserved in ≥99% of all species. Our analysis reveals that the most evolutionarily rigid components of the eukaryotic cell are the structural and maintenance machineries (**Figure 6**). The 99% Strict Core is heavily enriched for cytoskeletal elements (e.g., microtubule-based process, 13.9%) and chaperones (e.g., stress response, 3.3%). This is visually exemplified by the near-perfect structural superposition of core proteins such as Actin, Tubulin, Histone H3, and HSP70 (Heat shock 70 kDa protein) between distant lineages (**Figure 6, right panels**).

**Figure 6.**
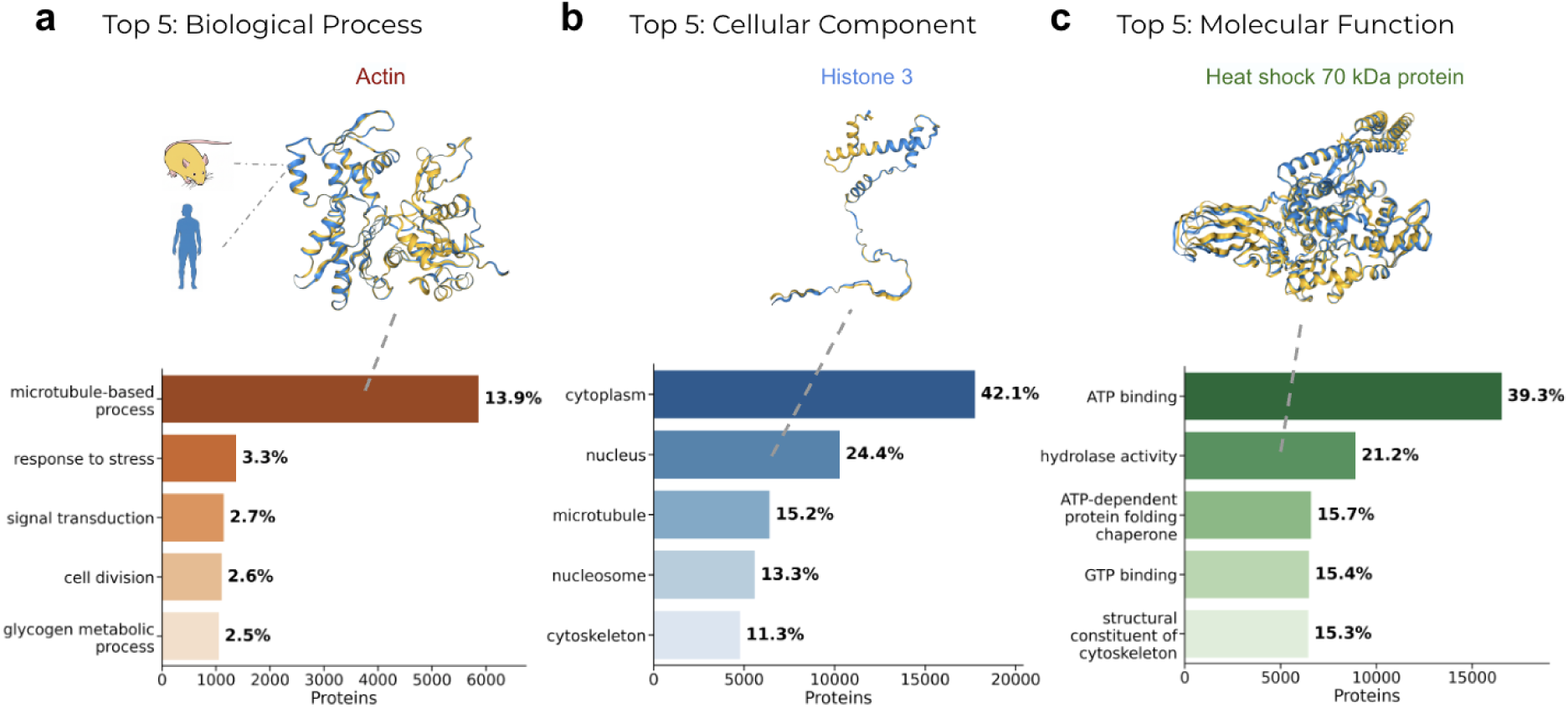
Identification and structural conservation of the core eukaryotic proteome. Gene Ontology (GO) frequency analysis was performed on the "Strict Core" protein families conserved in ≥99% of all analysed species. Bars represent the absolute number of proteins annotated with each term, with percentages indicating relative abundance within the total core dataset. Representative structural alignments between Human (blue) and Mouse (yellow) orthologs are displayed above their corresponding categories to illustrate the extreme architectural preservation of the strict core. Key examples include Actin (a), Histone 3 (b), and the Heat shock 70 kDa protein (HSP70, c). **a)** Top 5 Biological Processes: The core is dominated by structural integrity and regulation, most notably microtubule-based processes (13.9%), rather than metabolic housekeeping. **b)** Top 5 Cellular Components: Results highlight the prominence of the cytoplasm (42.1%), nucleus (24.4%), and cytoskeleton (11.3%) as the conserved spatial framework of eukaryotic cells. **c)** Top 5 Molecular Functions: Analysis reveals a heavy reliance on energy consumption (ATP/GTP binding) and structural quality control, such as ATP-dependent protein folding chaperones.

Interestingly, as we relaxed the conservation threshold to include families present in ≥95%, ≥90%, and ≥85% of species (**Supplementary Figure 1**), the functional landscape shifted. Unlike the strict core, these broader sets are dominated by translational machinery. At the 90% threshold (comprising ∼2 million proteins), "translation" becomes the top biological process (5.1%), and ribosome-related functions appear frequently. This suggests that while the translational apparatus is universally essential, individual components exhibit slight evolutionary turnover or loss in rare lineages, excluding them from the strict 99% set. Conversely, the cytoskeletal and chaperone networks identified in the Strict Core represent the immutable structural foundation of eukaryotic life.

### SHE as a Dual Indicator of Evolution and Annotation

To benchmark SHE against classical markers, we compared primate phylogenies derived from 18S rRNA sequences versus our proteome-wide structural approach **(Figures 7a&b**). The SHE-derived tree demonstrates superior statistical stability, yielding consistently high bootstrap values (71–98%) compared to the variable support observed in the rRNA tree (21–100%), which often suffers from incomplete sequence sampling (e.g., missing *Mandrillus leucophaeus* data).

**Figure 7.**
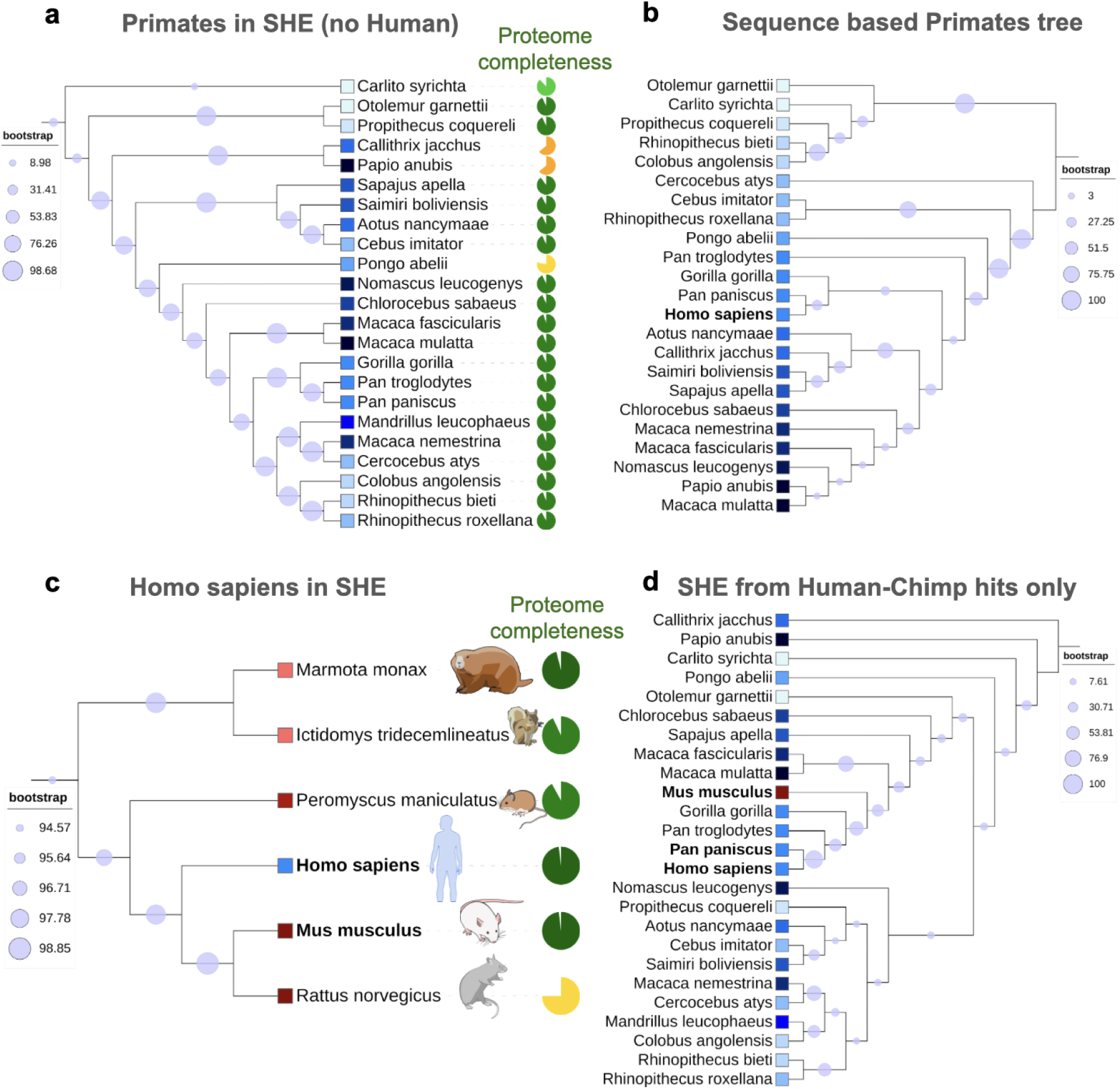
Proteome Completeness and the Structural Phylogeny of Primates. Structural phylogeny reveals that annotation maturity can influence local topology, serving as a diagnostic signal for database completeness. **a) Primate Phylogeny in SHE:** A subtree extracted directly from the global SHE structural phylogeny. While bootstrap support values (violet) remain high, the topology is influenced by the proteome completeness of each species (represented by pie charts), highlighting the relationship between available structural data and branching order. **b) 18S rRNA Reference Tree:** A benchmark phylogeny constructed from 18S rRNA sequences. Bootstrap support values (violet) show less consistency compared to the structural SHE tree. Species are color-coded by a shade of blue to facilitate a direct comparison of clustering order between sequence-based and structure-based methods. **c) Structural Similarity vs. Taxonomic Divergence:** A detailed subtree illustrating the anomalous clustering of *Homo sapiens* (human) with rodent species. Both humans and the house mouse (*Mus musculus*) feature high proteome completeness (∼98%) and share 16,513 structural orthologs. The combination of high aggregate bit-scores and near-complete structural coverage drives their close proximity in the global tree. **d) Normalisation of Annotation Bias:** A SHE phylogeny pruned to a uniform set of 15,186 structural hits shared between humans and *Pan paniscus* (bonobo). By removing the impact of missing data in primate annotations, *Homo sapiens* correctly clusters with its closest relatives, including *Pan troglodytes* (chimpanzee) and *Gorilla gorilla*. The inclusion of *Mus musculus* (red) as an outgroup demonstrates that when annotation gaps are eliminated, structural distance accurately reflects evolutionary divergence.

However, while SHE provides a robust global framework, it reveals a topological discrepancy in the placement of *Homo sapiens*. Contrary to established taxonomy, which groups humans with *Pan* (bonobos and chimpanzees) and *Gorilla* [25], the global SHE tree clusters humans more closely with highly annotated rodents (**Figure 7c**). Specifically, the prairie deer mouse (*Peromyscus maniculatus*, distance 0.47) and house mouse (*Mus musculus*, 0.49) appear as humans’ closest relatives, while chimpanzees appear significantly more distant (9.52).

We identify this discrepancy not as a biological signal, but as a measure of annotation maturity. Because SHE calculates distance based on cumulative alignment scores, missing proteins in less-annotated species act as "evolutionary penalties." The *Homo sapiens* and *Mus musculus* proteomes are nearly complete (>98% coverage in UniProt+AFDB), whereas the *Pan troglodytes* proteome is only ∼93.9% complete, and *Papio anubis* drops to ∼64% (**Figure 7a**). Consequently, humans and mice accumulate the highest density of high-scoring structural pairs, driving their artificial clustering.

To validate this mechanism, we reconstructed the primate tree using restricted protein subsets. When the input was pruned to include *only* the 15,186 proteins shared between *Homo sapiens* and *Pan paniscus*, humans correctly clustered with bonobos and chimpanzees (**Figure 7d**). Similarly, utilising the 12,310 proteins shared with *Papio anubis* corrected the human placement relative to baboons (**Supplementary Figure 3a**). In these controlled analyses, *Mus musculus* shifted to its correct outgroup position. This is especially evident in the tree pruned to the 2,773 core proteins common to all Primates (**Supplementary Figure 3b**). This confirms that SHE functions as a dual-purpose tool: it measures evolutionary structural distance while simultaneously serving as a sensitive detector of gaps in global proteome annotation.

### The SHE Web Server

To democratize access to the nearly 300 trillion (2.96 × 10^14^ ) structural alignments generated in this study, we have developed a comprehensive, freely accessible web platform: https://she-app.serve.scilifelab.se/. The application transforms the static data of the SHE project into a dynamic workspace for evolutionary analysis, enabling researchers to explore the structural history of eukaryotes without requiring massive local computational resources. A core feature of the server is the "Model Organism Discovery" module, which streamlines the search for optimal experimental models. By inputting any protein or pathway of interest, facilitated by guided examples, users can calculate structural conservation scores across potential model species. These rankings are derived from cumulative Foldseek bitscores and are accompanied by direct visualisations of the aligned protein structures, allowing for rapid assessment of structural fidelity.

Beyond specific pathway analysis, the platform offers a powerful environment for interactive phylogenetic exploration. The global SHE tree, encompassing 1,542 species, can be navigated in real-time, allowing users to zoom into major superkingdoms such as Opisthokonta, SAR, and Viridiplantae (**Figure 8**). The visualisation interface supports multiple layouts, including circular and linear dendrograms, and allows the tree to be dynamically colored by evolutionary rate or by the presence/absence of specific structural homologs. This functionality enables researchers to visually track gene loss events or identify evolutionary accelerators within distinct clades. Finally, to support further computational innovation, all underlying resources are made available for bulk download, including the complete matrix of Foldseek bitscores, the compressed Foldcomp databases of AlphaFold v4 predicted structures, and the Newick files for the global SHE tree and all extracted subtrees.

**Figure 8.**
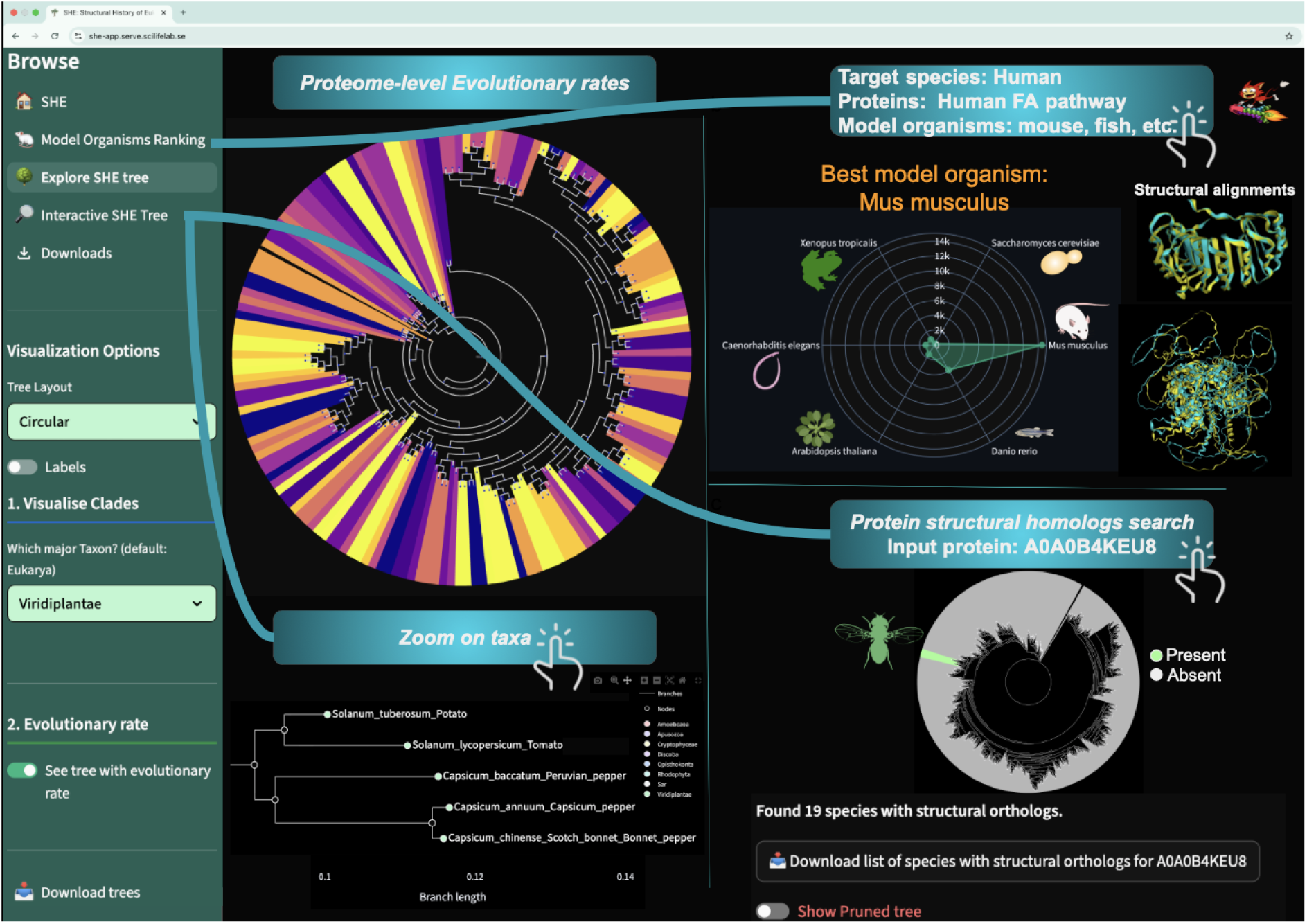
The SHE Web Server Interface. The Streamlit-based application enables real-time interaction with the structural phylogenetic data. **a)** Interactive Phylogenetic Exploration: The SHE app enables real-time visualisation of the eukaryotic tree of life. Users can toggle visualisation modes, such as colouring branches by Evolutionary Rate, and zoom into specific sub-kingdoms or species. Tooltips provide immediate access to proteome statistics, including size and calculated divergence rates. **b)** Data-Driven Model Organism Ranking: Demonstrating the power of proteome-wide structural matching, the "Model Organism" tool accepts a set of human proteins (e.g., the FA pathway) and ranks potential model species based on structural conservation scores. This allows researchers to quantitatively validate experimental models, distinguishing between high-conservation species (e.g., *Mus musculus*) and divergent lineages. **c)** Structural Homology and Tree Pruning: The function maps the presence/absence of structural homologs for any query protein. The interface automatically generates a "pruned" phylogeny (right), displaying the evolutionary trajectory of specific families, such as the lineage-restricted FXNA-like protease in *Drosophila* or universally conserved "housekeeping" proteins.

## Discussion

We present a high-resolution phylogenetic resource derived from the all-vs-all comparison of 1,542 eukaryotic proteomes. By converging rapid structural alignment [11] with mass-scale prediction [9], this work transcends the computational barriers that previously limited structural phylogenetics to small datasets. Leveraging the speed of Foldseek and the compression of Foldcomp [12], we integrated nearly 300 trillion structural alignments, demonstrating that proteome-scale analysis is not only feasible but essential for resolving the global evolutionary landscape beyond isolated genetic markers.

Structural phylogenetics provides a dual advantage: the stability of protein folds detects deep evolutionary signals, while whole-proteome integration ensures robust statistical support (**Figure 7**). Beyond topology, our R_proteome_ metric reveals lineage-specific accelerations, most notably within Class Aves, proving that these rates are intrinsic biological signals rather than artefacts of proteome size (**Figure 5a**).

Furthermore, we resolve the eukaryotic proteome into two distinct evolutionary signatures. Challenging sequence-based inferences of a metabolically complex LECA [24], structural analysis defines the "Strict Core" (≥99%) as primarily architectural, dominated by cytoskeletal elements and chaperones (**Figure 6**). As conservation relaxes to 85–95%, the landscape shifts to an "Operational Engine" driven by translational machinery. This suggests a bipartite model of eukaryotic evolution: a rigid structural scaffold supporting a plastic, yet universally essential, protein synthesis apparatus.

Selecting the right model organism is often the rate-limiting step in translational research. SHE transforms this process from a heuristic approach into a quantitative, data-driven query **(Figure 2)**. By pre-computing conservation scores for millions of protein pairs, our platform allows researchers to empirically rank model species based on structural fidelity. As demonstrated with the Fanconi Anaemia pathway, this capability validates established models while uncovering non-traditional candidates (e.g., *C. elegans* for specific sub-modules), thereby optimising experimental design and reducing the risk of translational failure.

A critical insight from this study is the sensitivity of proteome-wide phylogenetics to data completeness. The anomalous clustering of *Homo sapiens* with *Mus musculus*, driven by their high annotation completeness (>98%) relative to the chimpanzee (∼94%), serves as a quantified measure of database maturity. Rather than a mere limitation, we propose this sensitivity as a diagnostic feature: SHE serves as a composite signal of evolutionary distance and annotation integrity.

These results demonstrate that while structural phylogenetics effectively captures deep evolutionary signals, species-level topology remains sensitive to proteome coverage. Consequently, SHE functions as a diagnostic benchmark for reference proteome quality, highlighting lineages where asymmetric annotation coverage, rather than biological divergence, alters local evolutionary trajectories. Future iterations may address these discrepancies through the coverage-based normalisation or subsampling strategies validated in **Figure 7**.

Furthermore, while the computational scale of this initial matrix required a Neighbour-Joining (NJ) approach, the future adaptation of Maximum Likelihood or Bayesian models to the structural domain will likely enhance fine-scale resolution. Nevertheless, the integrity of the global phylogeny is evidenced by the robust recovery of kingdom-level relationships (**Figure 1**). This macro-level stability confirms that a dominant structural evolutionary signal persists despite localised annotation noise; had the clustering been a mere artefact of data volume, these well-established biological hierarchies would be entirely obscured.

The future of high-resolution phylogenetics lies in the integration of structural data. As the coverage of the eukaryotic tree expands and proteome annotations improve, we anticipate that resources like SHE will redefine our understanding of deep evolutionary relationships. The framework presented here is designed for continuous growth, ready to ingest the next generation of structural predictions. By making these tools and data freely available, SHE provides a foundational platform for the community to explore the complex history of life, from the architecture of the first eukaryote to the molecular mechanisms of modern disease.

## Methods

### Data

#### UniProt

We selected all eukaryotic reference proteomes from UniProt [14] on August 9 2023 (n=2248). We then select only the non-redundant ones (one proteome per organism ID) and have ≤20% missing genes according to BUSCO (The Benchmarking Universal Single-Copy Ortholog assessment tool), resulting in 1803 proteomes. All genes were downloaded in fasta format for all proteomes, and the UniProt identifiers were extracted from the fasta files. However, not all proteomes had all UniProt identifiers present in the AlphaFold DataBase (AFDB, uniprot_v4) [10]. The median retained fraction is 95.5%, and we selected the proteomes that had at least 25% of entries in the AFDB (see Foldcomp), resulting in 1542 proteomes (86%) in the final selection.

The median proteome size is 13,898 proteins (**Figure 9a**). The smallest proteome (*Ordospora colligata*, a gut parasite in shrimp) has 1810 genes, and the biggest (*Araneus ventricosus*, Orbweaver spider) has 246,021 genes. In the taxonomic prevalence (**Figure 9b**), Opisthokonta is by far the most prevalent (1243 proteomes), followed by Viridiplantae (176 proteomes) and SAR (96 proteomes). The strong bias to Opisthokonta demonstrates our preference to study species related to our own and how understudied the other groupings are. This can also be seen in the proteome size distribution, where it seems like proteomes with more than 50,000 proteins are lacking.

**Figure 9.**
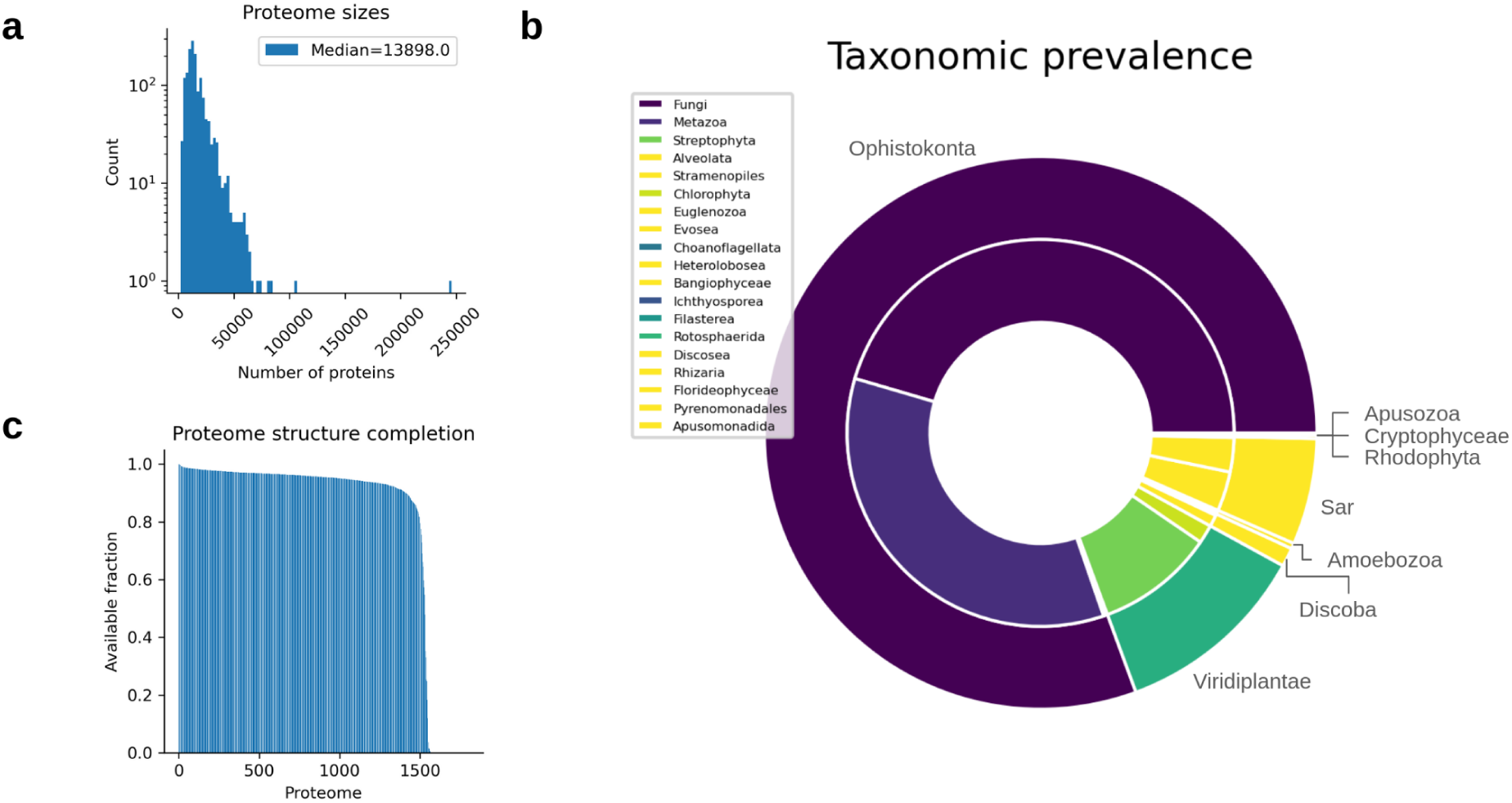
**a)** Distribution of proteome sizes. **b)** Fractions of proteins per kingdom (text on circle) and subkingdom (list). The Ophistokonta are the most represented Eukarya, followed by plants (Viridiplantae) and SAR. Among the Ophistokonta, fungi are the most prevalent, followed by Metazoa (animals). **c)** Available fractions of proteomes in the AFDB. Most proteomes have >90% availability.

#### Foldcomp

To save space and speed up the pairwise comparisons, we downloaded a compressed version of all predicted proteins from the AFDB uniprot_v4 [10] from Foldcomp (https://foldcomp.steineggerlab.workers.dev/, afdb_uniprot_v4). This database contains 214’683’829 protein structures and is 1.1 TB. We then extracted subsets of the compressed database according to the UniProt IDs from each proteome specified above. We used mmseqs2 (version ad6dfc66d7bbc4fd626fc19adf10ba587bc137c4) and the following command:

~~~
mmseqs createsubdb --subdb-mode 0 --id-mode 1 proteome_ids
input_foldcomp_db output_foldcomp_db
~~~

From 1803 proteomes, 1542 could be subset with at least 25% retained proteins (median retention=95.5%, **Figure 9c**). This is because the gene identifiers and proteomes are constantly updated in UniProt, and the AlphaFold predictions are of UniProt version 2021_04. In addition, proteins with fewer than 16 or more than 2700 residues are excluded from the AlphaFold database.

#### FoldSeek Databases

From the proteome-specific databases created with Foldcomp and mmseqs2, we build FoldSeek (version d8fab94f544fa5014274a376344c2cfd7d7530bd) databases for structural alignment using the following command:

~~~
foldseek createdb folcomp_db foldseek_db
~~~

### Protein Structural Similarity Search with FoldSeek

To compare all proteins between entire proteomes, we use FoldSeek and the built databases of all structures in each proteome (see above) to be able to search these efficiently. FoldSeek computes a structural bit score that combines the 3Di alignment and structural scores: “*We rank hits by a ‘structural bit’ score - that is, the product of the bit score produced by the Smith-Waterman algorithm and the geometric mean of average alignment LDDT and the alignment TM-score.” [11]*

For each proteome i,.., n-1:

1. Search proteome i+1..n with FoldSeek, where n denotes the total number of proteomes. The following command was used for the search:

~~~
foldseek search foldseek_db(i) foldseek_db(i+1) aln(i,i+1)
<tmpDir> -e 0.01
~~~

2. Thereafter, the results (with evalue<0.01) are formatted into a tsv:

~~~
Foldseek convertalis foldseek_db(i) foldseek_db(i+1) aln(i,i+1)
result(i,i+1)--format-output "query,target,alnlen,evalue,bits"
~~~

3. The highest bit score for each protein in proteome i is selected as the top hit in proteome i+1. This ensures the proteins’ structures are similar. Each protein in proteome i+1 is used only once. This means that if two proteins have identical top hits, one will get a score of 0. This ensures differences between duplicated genomes, which are ignored by conventional phylogenetic methods.

Searching 1542 proteomes against each other (excluding self-interactions) results in (1542×(1542-1))/2 = 1,188,111 comparisons. Applying the same calculation to the number of proteins in each proteome results in 2.96 × 10^14^ protein alignments between the 24,333,776 unique proteins.

### Proteome-wide Structural Distance Calculations

To calculate the distance between two proteomes, we use all obtained top hit bit scores for all comparisons in the protein similarity calculations (n=2.96 × 10^14^). **Figure 10a** shows the distribution of all bit scores, and equations 1 and 2 in the main text specify the proteome similarity and distance, respectively. We sort the proteomes by the number of proteins and search the biggest against the smallest to capture size differences (bit score=0 for missing matches). A distance matrix is produced by calculating all pairwise distances between all proteomes (**Figure 10b**).

**Figure 10.**
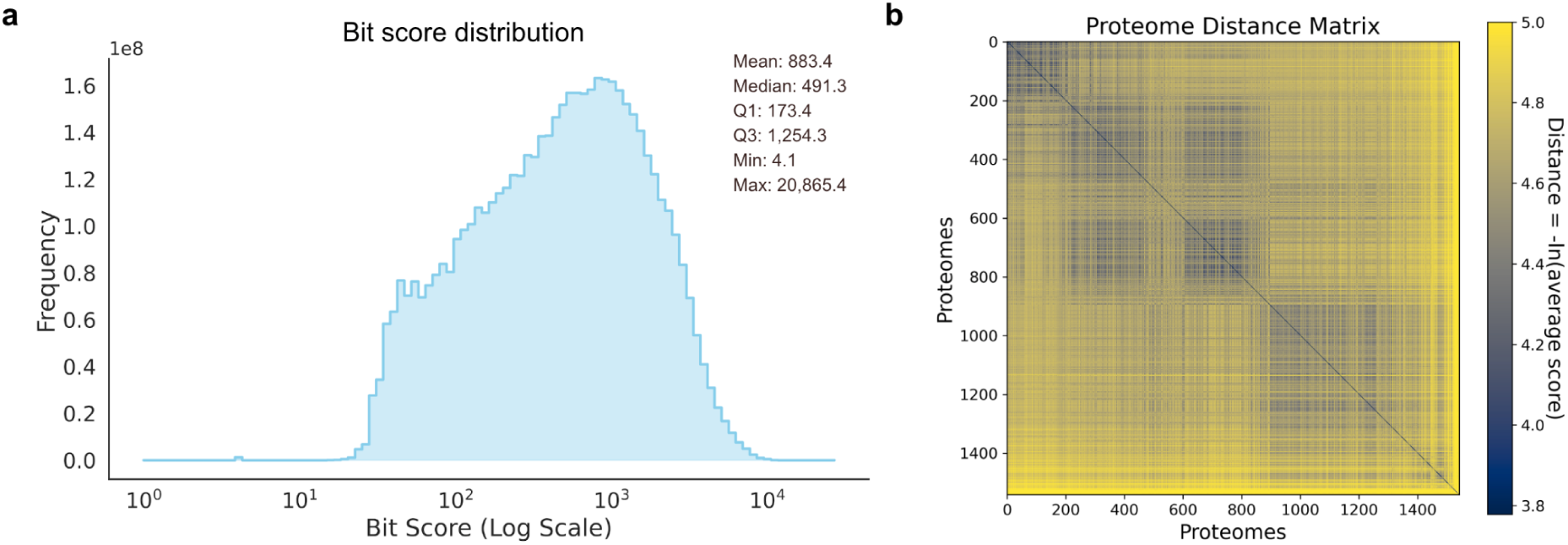
Global analysis of proteome similarity and distance. **a)** Histogram and corresponding Kernel Density Estimate (KDE) showing the frequency distribution of summed bit scores from all pairwise proteome comparisons. The x-axis shows the base-10 logarithm of the bit score sum. This is a measure of overall sequence similarity between two proteomes. The y-axis indicates the count of proteome pairs for each bit score sum. **b)** Heatmap of the pairwise evolutionary distances between proteomes. The distance, indicated by the colour scale, is calculated as the negative natural logarithm (-ln) of the mean similarity score. Darker colours represent smaller distances and higher similarity, while lighter colours indicate larger distances and lower similarity. The diagonal results from the self-comparison of each proteome, having a distance of zero.

### Neighbour Joining

Unlike sequence-based phylogenetics, which models discrete substitution processes, our approach operates on continuous structural similarity aggregated across complete proteomes and therefore does not assume an explicit evolutionary substitution model. We use neighbour-joining (NJ [26]) following FoldTree [5] to construct the evolutionary relationship between all proteomes. This method produces accurate relationships given that the distances between the proteomes are accurate [27]. More advanced methods, such as maximum likelihood or parsimony, exist but are not possible to apply on this scale, and it is not known what model of evolution is the most accurate. Parsimony methods assume that the smallest, simplest trees are the best, but it is possible that evolution does not work in this way at all. Instead, we choose to rely on the overall structural similarity between two proteomes and NJ [26].

### Bootstrapping

A phylogenetic tree is usually bootstrapped to build a consensus tree and to obtain estimates of the confidence in the branching [28]. As it is too computationally demanding to recompute alignments and no evidence-based model for structural substitutions in evolution exists (to our knowledge), we apply this methodology to the distance matrix directly. By randomly removing pairwise relationships, we build a tree using 10,000 replicates with the Phylo package from BioPython (version 1.81). We employed 10,000 replicates to ensure topological stability, as lower counts produced unresolved polytomies. While the count of strongly supported nodes (≥70%) converged early, the total resolution continued to improve with replicate count (**Supplementary Figure 2a, Supplementary Table 1**). Specifically, 10,000 replicates resolved 1,539 internal nodes, which is close to the theoretical maximum of 1,541. In contrast, 1,000 replicates yielded only 1,525 nodes. Median bootstrap support remained consistent (∼89%) across replicates, while minimum support values converged near zero, indicating the stable recovery of distinct low-support nodes (**Supplementary Figure 2b, Supplementary Table 2).**

### Normalisation of the Phylogenetic tree

To improve the visual clarity of the phylogenetic tree, all branch lengths were subjected to a two-step mathematical transformation. First, the lengths were scaled to a range between 0 and 1 using min-max normalisation.

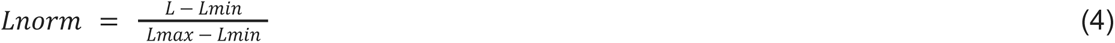

Here, L_min_ and L_max_ are the minimum and maximum raw branch lengths in the tree, respectively. Second, to invert the scale and expand the differences between the shortest branches, a negative logarithmic transformation was applied to the normalised lengths to compute the final branch length, L_new_.

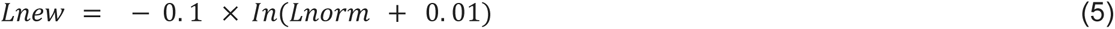

This procedure was designed to enhance the visual spread between clades, with the constant 0.01 serving to avoid an undefined result for zero-length branches. The resulting phylogeny retains its original topology, but features rescaled branch lengths optimised for clearer interpretation.

### Calculating the Structural Evolution of Protein Sets

#### Model Organism Selection

To identify the most suitable model organisms for the study of the FANCD2-dependent DNA repair pathway, we developed a scoring pipeline. A list of target human UniProt identifiers (for example: Q9BXW9, Q9NXK8, O14757, Q9NVI1, Q13616) was queried against pairwise proteome alignment datasets comparing Homo sapiens to a panel of candidate model organisms (**Figure 2**). For each candidate species, we calculated a cumulative bit score by summing the maximum bit scores of the identified orthologs *(equation 1)*. To account for bidirectional database orientation, the pipeline automatically detected and processed both forward (Target-vs-Organism) and reverse (Organism-vs-Target) alignment files.

#### Family-specific Phylogeny

To extract the family-specific phylogenies, we set a strict bit score cutoff of ≥ 173 (corresponding to an E-value of 0), which aligns with the first quartile (Q1) of the bit score distribution (**Figure 10a**) and effectively retains the top 75% of the most significant structural alignments. We then created a similarity graph. In this graph, each protein is a node, and an edge indicates a statistically significant structural alignment. We divided the graph into distinct protein families using a connected components algorithm from the Python library NetworkX [29]. Each resulting component represents a group of structurally related orthologs and paralogs. Finally, we utilised the ETE3 toolkit [30] to generate the phylogeny for each isolated family (**Figure 3**). We mapped the species IDs of the proteins within each family to their corresponding organism names in the global tree. We then pruned the global consensus tree to retain only the leaves corresponding to the species present in that specific family, preserving the original branch lengths.

### Evolutionary Rate Calculations

To quantify the tempo of structural divergence across the eukaryotic tree, we calculated a proteome-wide evolutionary rate (R_proteome_). This metric represents the ratio of structural change in a given lineage relative to its immediate ancestor, as defined in Equation 3 (see Results, **Figure 4**).

The rates were calculated by traversing the Neighbour-Joining tree toward the terminal nodes. For each node, the "current" branch length was compared to the "parental" branch length. To ensure the robustness of the reported accelerations and decelerations, we implemented a numerical stability filter: if the ancestral branch length (the denominator) was less than 10^-3^ structural distance units, the rate was excluded from the analysis (assigned as 0 and not used for inferring evolutionary rates, effectively NA).

This thresholding was necessary because branch lengths nearing zero indicate that the lineages are too structurally similar to calculate a meaningful ratio, and excluding them correctly identifies them as areas of the tree where the structural signal is below the resolution limit for rate comparison. All subsequent distributions and outlier analyses (e.g., the lineage-specific accelerations in *Class Aves*) were performed using this filtered dataset to ensure that high R_proteome_ values reflect genuine biological shifts rather than computational artefacts.

### Proteome-Scale Data and Phylogenetic Resolution Calculations

Traditional phylogenomics relies on substitution models (e.g., Jukes-Cantor [31]) that assume discrete character states in DNA or amino acid sequences. However, these models do not apply to structural proteomics, where evolution is measured through continuous geometric similarity and fold conservation rather than point mutations. Because structural folds are more conserved than sequences, they can persist over vast evolutionary timescales, but they also risk signal "saturation" or noise when analysing entire proteomes.

We hypothesised that the aggregate structural signal from thousands of protein families would provide a more stable and higher-resolution evolutionary map than isolated markers. To test this and determine the minimum data threshold required for a robust structural phylogeny, we conducted an experiment utilising five nested subsets of widely distributed orthologous families.

These subsets were designed with increasing scales of structural data: Level 1 (6,989 proteins, 1 family), Level 2 (12,006 proteins, 2 families), Level 3 (101,674 proteins, 39 families), Level 4 (1,000,112 proteins, 2,362 families), and Level 5 (2,176,597 proteins, 178,220 families) **(Figure 5b)**. For each subset, we generated a pairwise inter-proteome distance matrix derived from the aggregate Foldseek bit scores of the included proteins. The distance between any two proteomes was calculated using ***Equation 2*** with the addition of a constant factor of 2 to the denominator to ensure mathematical stability at the lower end of the bit-score spectrum. Using these matrices, we constructed phylogenetic trees via the Neighbour-Joining (NJ) algorithm and evaluated nodal confidence through 10,000 bootstrap replicates. This iterative scaling allowed us to quantify the gain in statistical support and taxonomic resolution provided by transitioning from single-family markers to proteome-wide structural datasets.

### The Core of Eukarya and GO Enrichment

To find a structural core shared by all Eukarya, we used a filtering strategy focused on structural similarities and taxonomic distribution. We compared all the proteomes in a complete structural analysis. First, we created a list of orthologous protein families from hits with a FoldSeek bit score over 173 and an E-value of 0.0005. Next, we grouped protein families generated via a graph clustering method, Markov Cluster Algorithm (MCL) [32]. We applied MCL to the weighted network of homologous protein pairs using an inflation parameter (I) of 2.0 to balance cluster granularity. To functionally characterise these families, we extracted all constituent protein IDs and mapped them to UniProtKB accessions using the UniProt ID Mapping API [14,33]. Gene Ontology (GO) [34] terms (Biological Process, Molecular Function, and Cellular Component) were retrieved for every protein within the identified core families using the g:Profiler service [35]. To generate a global functional profile, we performed a frequency analysis of GO terms across the entire dataset for each threshold (99%, 95%, 90%, and 85%) (**Figure 6 and Supplementary Figure 1)**.

### Primate Phylogenetic Tree Construction

We compared the phylogeny of primates between the structure-based approach (created like above) and a sequence-based approach. We applied the ETE3 toolkit [30] to extract the phylogeny of primates from SHE (**Figure 7a**). To construct the sequence-based phylogenetic tree of primates (Figure 7b), we retrieved 24 primate 18S rRNA sequences from the NCBI nucleotide database [14,33]. Sequences were aligned using the MUSCLE [36]. Pairwise distances were then calculated using the Kimura 2-parameter (K2P) model [37] to generate a distance matrix suitable for 18S rRNA sequences, as it accounts for the higher frequency of transitions relative to transversions in ribosomal RNA. Phylogenetic analysis was performed using the NJ method with 10,000 bootstrap replicates to construct the tree and assess branch support. All alignments, distance calculations, and tree construction were conducted in MEGA 12 software [38]. To reconstruct phylogenetic evolution from a protein subset (i.e., **Figure 7c and Supplementary Figure 3**), a similar approach to the Model Organism selection was applied. We use a dataset of proteins as a reference query set (e.g. the hits between Homo sapiens and Pan Paniscus). For each comparison species (i.e. all Primates), we generated a structural similarity profile by extracting the maximum bit score for each input protein against all the comparison proteomes. The resulting score vectors for all species were aggregated into a distance matrix, which was further used to construct a phylogenetic tree using Neighbour Joining with 10,000 bootstraps.

## Acknowledgements

We thank DeepMind for the creation and sharing of AlphaFold2 and the AlphaFold database. We thank the creators of mmseqs2, FoldSeek and Foldcomp for their wonderful software. The ecosystem created by the Steinegger group for protein sequence and structural comparisons has made this work possible, and we foresee a new wave of evolutionary studies at a large scale due to these achievements. We thank the Interactive Tree Of Life (iTOL) v7 [39], an invaluable online tool for the display and annotation of the large-scale phylogenetic trees presented in this work. NIH Bioart was used to visualise the different Eukarya on the organism level. The SHE app was created using the Streamlit open source Python framework, and made freely accessible through SciLifeLab serve, for which we thank. We thank M. Madan Babu for useful discussions and the suggestion of the model organism matching.

## Contributions

PB designed the study, developed the workflow for proteome comparison and performed the proteome alignments. QL, together with PB and DD, built the tree from the resulting alignments and performed all subsequent analysis. DD made the web server. All authors contributed to writing and visualising the results. PB obtained funding.

## Availability

SHE and all its data are available at: https://she-app.serve.scilifelab.se/

## Supplementary Information

### Supplementary Figures

**Supplementary Figure 1.**
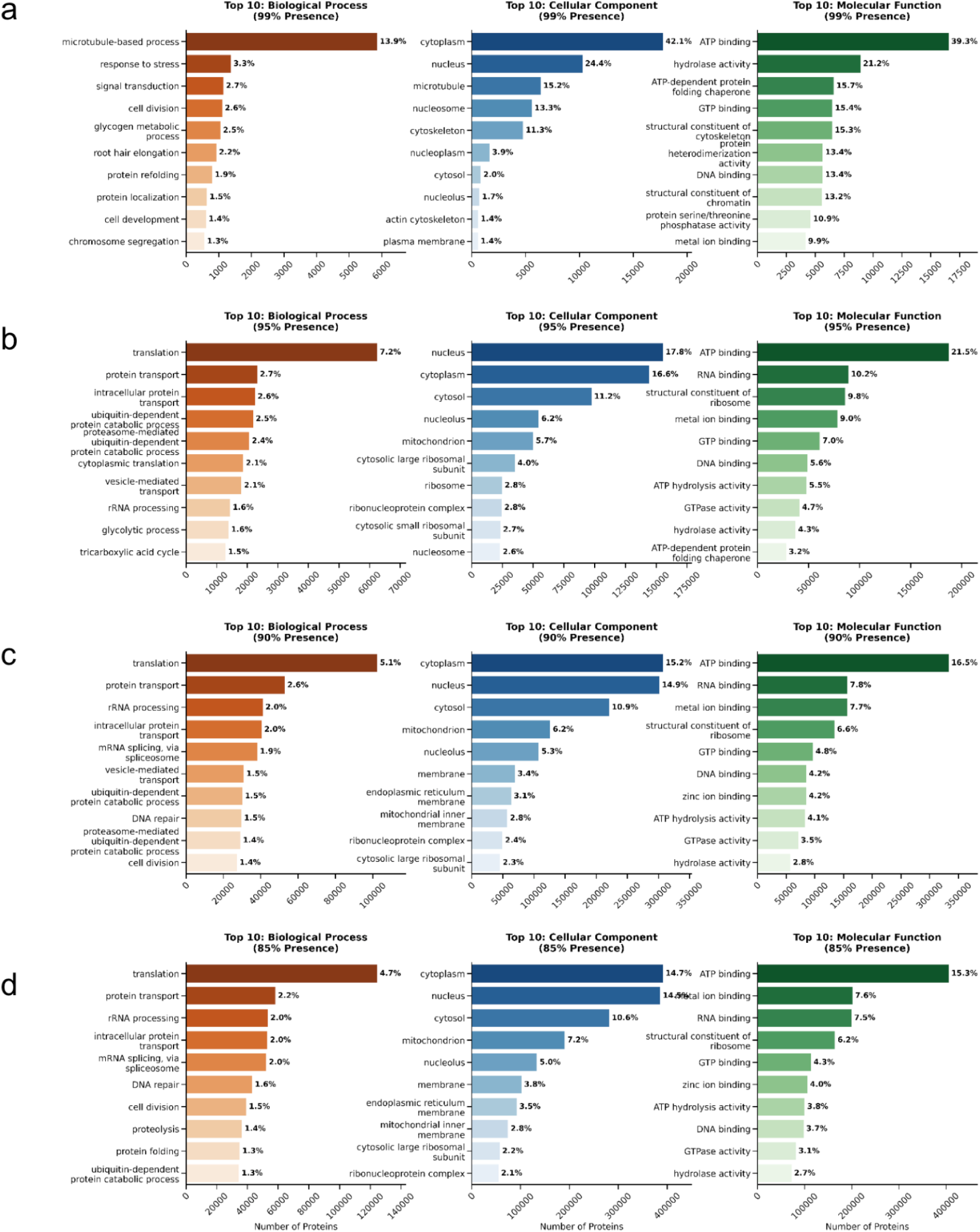
Functional Stratification and Scaling of the Eukaryotic Structural Core. Gene Ontology (GO) enrichment analysis of human non-intraspecies paralogous proteins across varying structural conservation thresholds. Bar lengths represent the absolute number of proteins annotated with each term, while the adjacent text labels indicate the percentage relative abundance of that term within the total core dataset for each specific threshold. **a)** The Strict Core (≥99% conservation; n = 42,128): Top 10 GO terms capture essential regulatory and architectural functions. Biological processes are dominated by cell division and protein refolding (chaperones), while kinase activities represent the primary molecular functions. **b)** The Intermediate Core (≥95% conservation; n = 874,787): At this relaxed threshold, the functional landscape shifts toward the emergence of translational machinery. Ribosomal subunits become the prevalent Cellular Components, contrasting with the cytoskeleton-dominated strict core in panel (a). **c)** The Operational Core (≥90% conservation; n = 2,022,223): This tier is dominated by universal housekeeping machinery. Translation (5.1%) emerges as the top Biological Process, and RNA binding rises in prominence due to the inclusion of diverse RNA-processing families. **d)** The Broad Core (≥85% conservation; n = 2,659,960): At the most relaxed threshold, the functional profile stabilises with increased volume. Translation (4.7%) and protein transport continue to define biological processes. ATP binding (15.3%) and metal ion binding are the dominant molecular functions, reflecting the broad enzymatic and cofactor requirements of the core proteome.

**Supplementary Figure 2:**
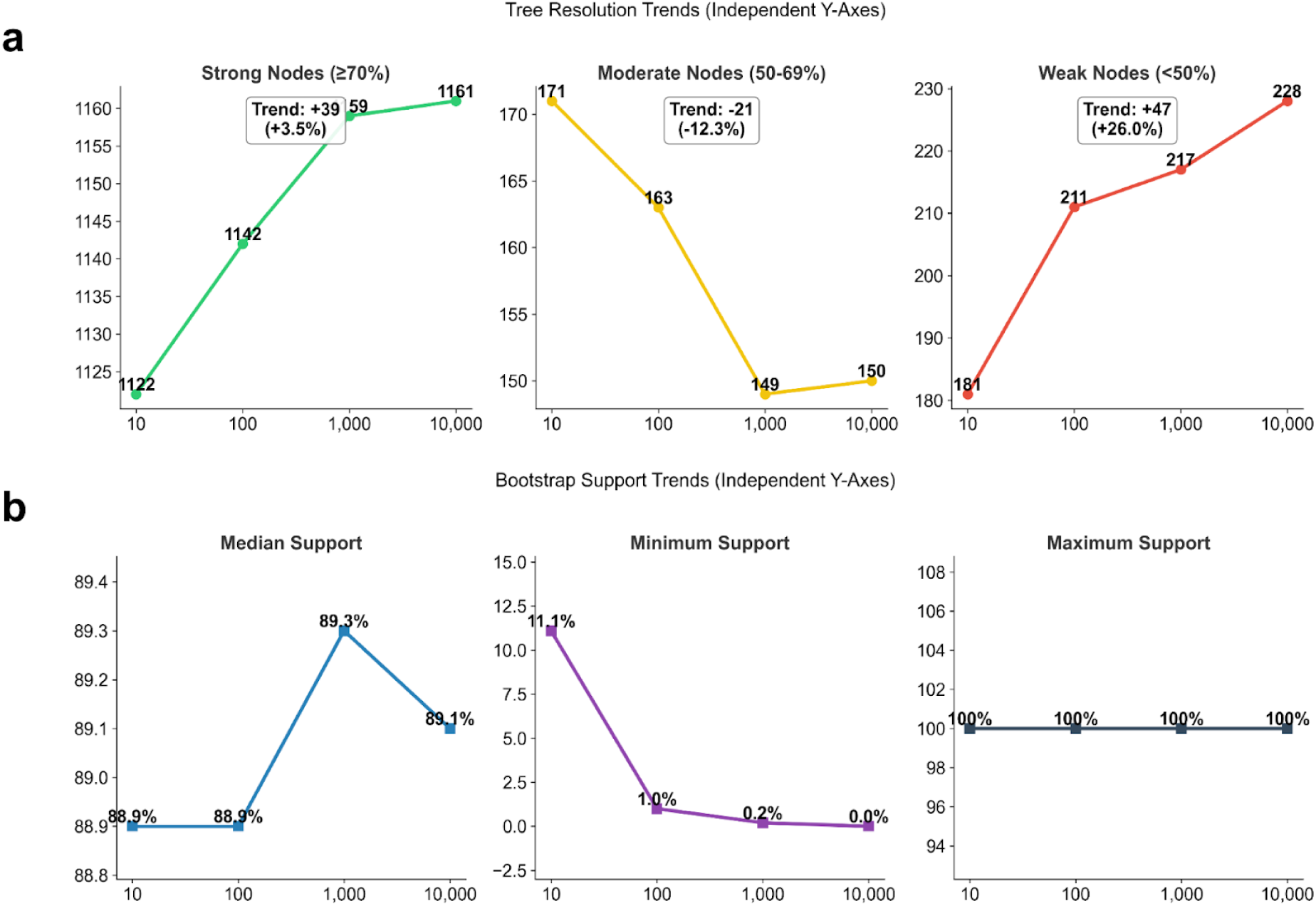
Convergence of bootstrap support values with an increasing number of replicates. The analysis was performed with 10, 100, 1,000, and 10,000 replicates. **a)** Trends in phylogenetic resolution categorised by node strength: Strong (≥ 70%), Moderate (50-69%), and Weak (< 50%). Panels are displayed with independent y-axes to highlight relative trends across different magnitudes. This view reveals that while strong nodes remain stable, there is a distinct increase in weak nodes and a decrease in moderate nodes, suggesting a shift from ambiguous to definitively unresolved states as sampling increases. **b)** Plot of key statistical metrics for bootstrap support across all nodes, showing that median and maximum support remain high while minimum support quickly converges to zero.

### Supplementary Tables

**Supplementary Table 1:** Classification of internal nodes by bootstrap support strength. Node counts are shown for 10, 100, 1,000, and 10,000 replicates. Nodes are stratified into Strong (≥70%), Moderate (50–69%), and Weak (<50%) categories. Total internal nodes indicate the degree of tree resolution relative to the theoretical maximum, demonstrating improved resolution.

| Metric | 10 Replicates | 100 Replicates | 1,000 Replicates | 10,000 Replicates |
| --- | --- | --- | --- | --- |
| Strong Nodes ($\geq 70\%$) | 1122 | 1142 | 1159 | 1161 |
| Moderate Nodes (50-69%) | 171 | 163 | 149 | 150 |
| Weak Nodes ($< 50\%$) | 181 | 211 | 217 | 228 |
| Total Internal Nodes | 1474 | 1516 | 1525 | 1539 |

**Supplementary Table 2:** Bootstrap supports value statistics across replicates. Median, minimum, and maximum support percentages calculated for 10, 100, 1,000, and 10,000 bootstrap replicates.

| Metric | 10 Replicates | 100 Replicates | 1,000 Replicates | 10,000 Replicates |
| --- | --- | --- | --- | --- |
| Median Support | 88.90% | 88.90% | 89.30% | 89.10% |
| Minimum Support | 11.10% | 1.00% | 0.20% | 0.00% |
| Maximum Support | 100.00% | 100.00% | 100.00% | 100.00% |

**Supplementary Figure 3.**
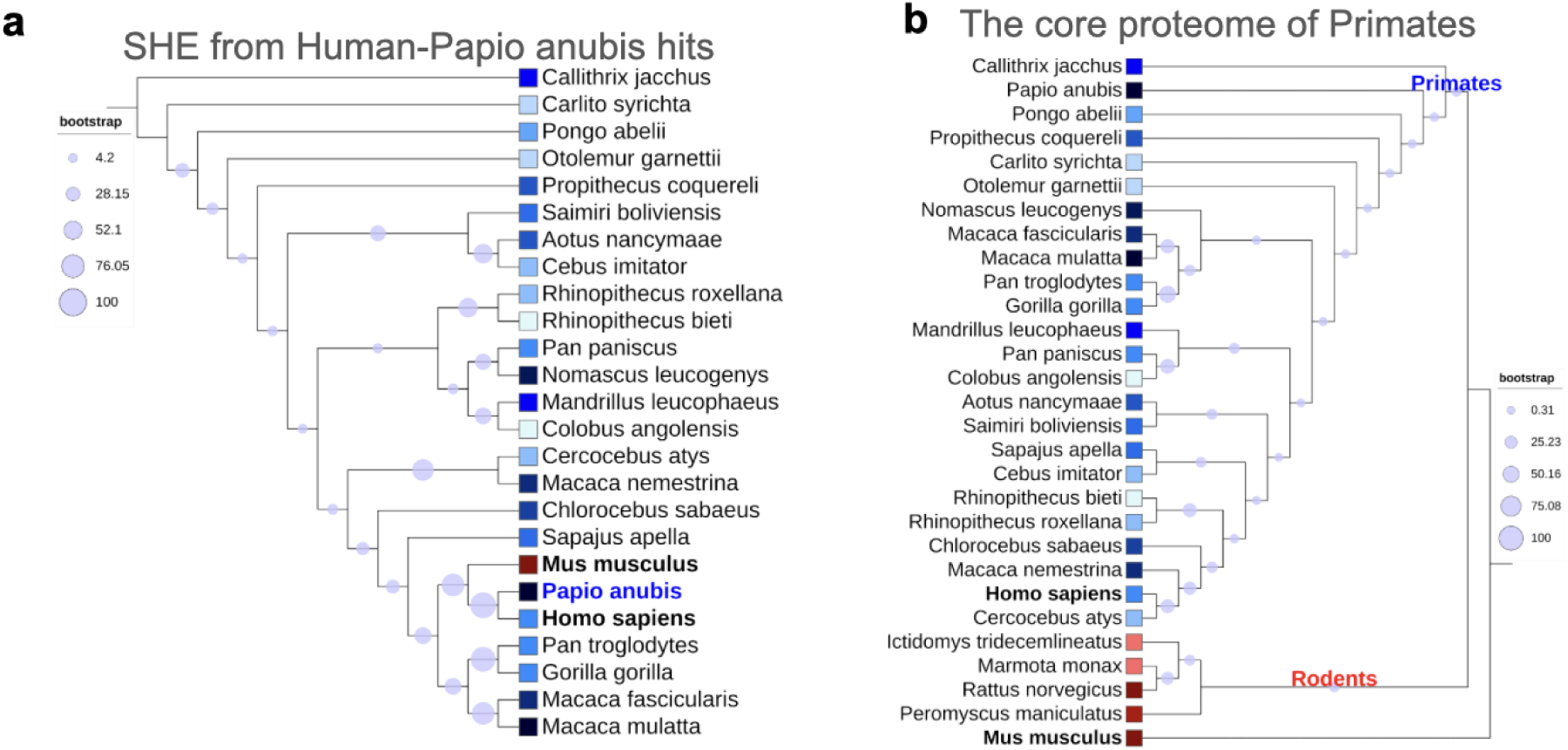
SHE, Primates and Rodents. **a)** SHE tree pruned to 12,310 Homo sapiens-Papio anubis (least proteome completeness among Primates - 64%) protein structural hits. The absence of missing hits drives the sampling of Human and Baboon together, confirming missing proteins as the driving factor in the construction of the tree. Here, the house mouse (Mus musculus) is included to further show the weight of no missing proteins in the tree-building method. **b)** SHE tree pruned to 2,773 core proteins among Primates (no missing proteins in Primates). Primates are thus clustered only based on the bitscore values from their structural comparisons. Rodents are now correctly clustered as an independent group because of the greater weight of missing proteins.

